# Potential of phenothiazines to synergistically block calmodulin and reactivate PP2A in cancer cells

**DOI:** 10.1101/2021.06.16.448716

**Authors:** Sunday Okutachi, Ganesh babu Manoharan, Daniel Abankwa

## Abstract

Phenothiazines (PTZ) are well known as inhibitors of monoamine neurotransmitter receptors, notably dopamine receptors. Because of this activity they are used for decades as antipsychotic drugs. In addition, significant anti-cancer properties have been ascribed to them. Several attempts for their repurposing were made, however, their incompletely understood polypharmacology is challenging.

Here we examined the potential of PTZ to synergistically act on two cancer associated targets, calmodulin (CaM) and the tumor suppressor protein phosphatase 2A (PP2A). Both proteins are known to modulate the Ras-MAPK pathway activity. Consistently, combinations of a CaM inhibitor and a PP2A activator synergistically inhibited cancer cells with KRAS or BRAF mutations. We identified the covalently reactive PTZ derivative fluphenazine mustard as an inhibitor of Ras driven proliferation and Ras membrane organization. We confirmed its anti-CaM activity in vitro and through a cellular CaM target engagement bioluminescence resonance energy transfer (BRET) assay. Our results suggest that improved PTZ derivatives retaining their synergistic CaM inhibitory and PP2A activating properties, but without neurological side-effects, may be interesting to pursue further as anti-cancer agents.

## Introduction

Several phenothiazines (PTZ) are approved as neuroleptics for the treatment e.g. of bipolar disorders. Structurally, they partially emulate dopamine, thus inhibiting dopamine D2 receptors, and affect also other monoamine neurotransmitter receptors (1). In addition, PTZ are known to inhibit calmodulin (CaM), the ubiquitous calcium binding protein, which regulates the activity of hundreds of proteins (2-4). CaM became a drug target in cancer, because of its profound involvement in regulating the cell cycle (3). Thus, attempts were made in the late 1980s to exploit the CaM inhibiting activity of PTZ in cancer therapy development (5). Such drug repurposing may offer a cheaper and faster access to live saving therapies. Since PTZ pass the blood brain barrier, the PTZ trifluoperazine was even tested in clinical trials to treat high grade gliomas (6).

Differential kinetic labelling experiments suggested that binding of trifluoperazine to Ca^2+^/CaM perturbs Lys75, 77 and 148, the same residues that are modified by the covalent CaM inhibitor ophiobolin A (7). In agreement with the kinetic labelling studies, chemically reactive PTZ modified the same lysines (8). Efforts were made to generate covalently binding derivatives of this compound class, as irreversible binding could significantly increase the potency of drugs (9). For instance, the PTZ fluphenazine (Flu), was functionalized with a chlor-ethylamine, also known as mustard-group, to generate a covalently reacting fluphenazine-mustard (Flu-M) derivative (10). Nitrogen mustards have been used for decades, because of their ability to crosslink the two strands of DNA (11). The mustard-group is converted into a very reactive intermediate aziridinium ion at neutral or basic pH by intramolecular nucleophilic substitution of the leaving chloride (11). The aziridinium then reacts with nearby nucleophiles, thus irreversibly linking the drug to the target, such as CaM.

While irreversible covalent inhibition can be regarded as the ultimate pharmacological inhibition strategy with potentially infinite ligand efficiencies, its exploitation in drug development was abandoned for many years. However, several successful drugs, such as aspirin and penicillin, rely on covalent inhibition mechanisms, and in recent years covalent inhibitor development has gained track again (9). One of the most promising developments in this area are the first direct inhibitors of K-Ras-G12C, which are equipped with an acrylamide electrophile to specifically react with the cysteine in the mutant form of K-Ras. These compounds bind to a cryptic pocket under the SII loop that is only accessible in GDP-bound K-Ras (12). Several realizations of such inhibitors exist from a number of companies, including the compounds ARS-1620, MRTX849 (Adagrasib) and AMG-510 (Sotorasib) (13-15). While Adagrasib is still in clinical trials, Sotorasib has been performing well in the clinic and was recently granted accelerated approval by the FDA for the treatment of KRAS-G12C mutant non-small cell lung cancers (16).

However, for other KRAS mutations and in other indications, there is still an urgent need for K-Ras inhibition strategies. Surrogate K-Ras targets, such as its trafficking chaperones CaM and PDE6D have emerged as interesting additional targets (17-19). The interplay between K-Ras4B (hereafter K-Ras) and CaM is particularly interesting, as it was implicated in the ability of K-Ras to drive cancer cell stemness (20). We previously showed that the natural product ophiobolin A (OphA), a covalent CaM inhibitor, can block stemness properties of cancer cells and K-Ras membrane organization (21). We recently developed less toxic functional analogues of OphA, which confirmed this potential (22). This lends an exciting new rational to the targeting of CaM, which justifies renewed interest in CaM inhibitor development (23).

Additional MAPK-signaling targeting strategies may act complementary or even synergistically. The tumor suppressor protein phosphatase 2A (PP2A) inactivates several oncogenic signaling pathways (24,25). Importantly, PP2A inactivation is required for the transformation of RAS mutant cancer cells, suggesting a particular importance of PP2A reactivation and Ras-pathway inhibition (26). PP2A actually comprises over 80 different holoenzymes, which are each composed of a scaffolding A subunit, a regulatory B subunit and the catalytic C subunit (27). The localization and specificity determining 15 distinct B subunits can be categorized into four families (28). The activity of the C subunit is critically regulated by leucine carboxmethyl transferase (LCMT), which reversibly attaches a methyl group to the C-terminal leucine to activate the phosphatase (29). This is counteracted by phosphatase methylesterase 1 (PME-1), which demethylates this site (25). Deregulation of the PP2A tumor suppressor in cancer typically occurs by the upregulation of endogenous inhibitors, such as cancerous inhibitor of PP2A (CIP2A), which selectively sequesters B56 family B subunits (30,31). Another endogenous inhibitor that is overexpressed for instance in lung cancer is SET (32).

Recently resistances against MEK-inhibition were shown to be overcome by novel PP2A activating drugs (33). These phosphatase activators, promise to modulate the onco-kinome more broadly than kinase inhibitors (34). These PP2A activators were derived from PTZ that were engineered for reduced central nervous system side-effects, such as sedation (35). Mechanistically, the small-molecule activator of PP2A (SMAP) DT-061 was shown to bind to an inter-subunit region formed by all three PP2A subunits, thus specifically stabilizing PP2A-B56α heterotrimers (36). The likewise PTZ derived iHAPs (improved heterocyclic activators of PP2A) function in the same way, however, slight differences in the B-subunit composition that is stabilized in the PP2A holoenzyme were found (37). It is not known, however, whether these compounds can still bind to CaM and as such may act on multiple targets.

Given that both CaM and PP2A are pharmacological targets of PTZ, it is possible that these drugs could combine within one molecule a potential synergistic activity of CaM inhibitors acting on the K-Ras stemness signaling axis and PP2A-activators suppressing the Ras transforming activity (21,33). Here we examined whether CaM inhibition and PP2A activation could synergize and if this synergistic activity can be re-unified in PTZ-like compounds or covalently reacting derivatives thereof. We propose that the synergistic polypharmacology of PTZ that is suggested by our data may explain some of their profound anti-tumor effects (38,39).

## Results

### Phenotypic assessment of CaM inhibition and PP2A activation reveal synergistic drug interaction commensurate with phenothiazine effects

We hypothesized that PTZ could synergistically combine the activities of CaM inhibition and PP2A activation. To assess this possibility, we first analyzed the combinatorial effect of the potent non-covalent inhibitor calmidazolium (CMZ) and the PP2A activating SMAP, DT-061, on the growth of KRAS mutant MDA-MB-231 breast and NCI-H358 lung cancer cell 3D spheroids, as well as BRAF mutant A375 skin cancer cell spheroids. These cell lines showed a spectrum of genetic dependencies on KRAS, HRAS, BRAF, CALM1-3 (the three human CaM genes), the alpha subunit of the PP2A enzyme (PPP2CA) and an endogenous inhibitor of PP2A in cancer, SET (**Figure 1A**).

**Figure 1.**
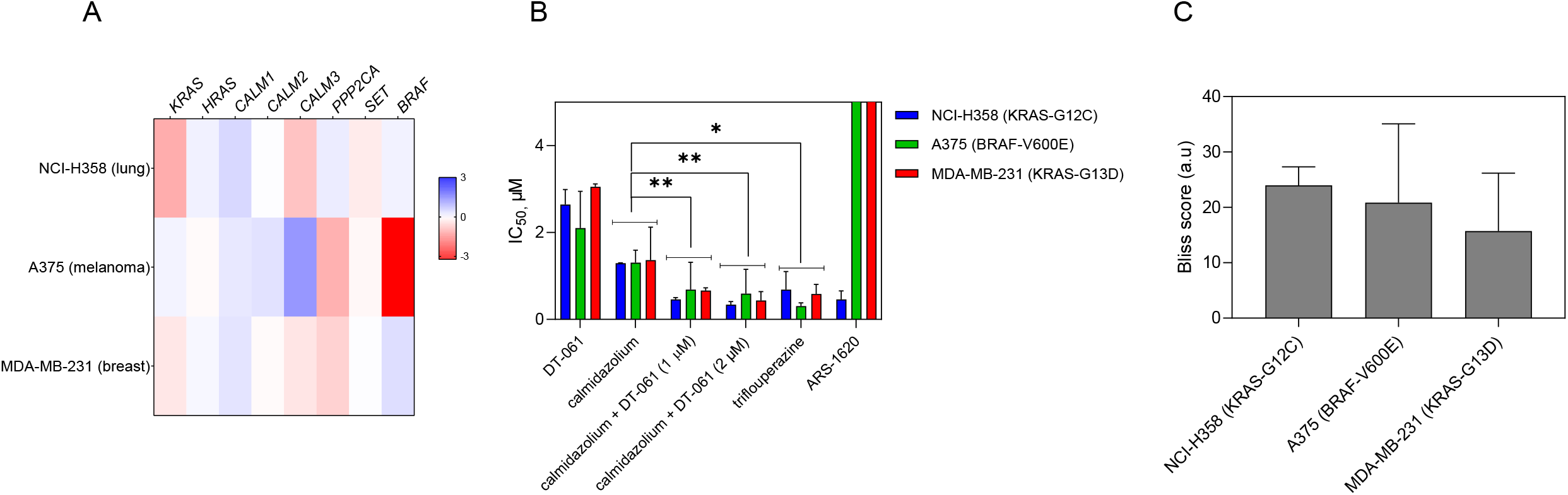
Cellular assessment of CaM inhibition and PP2A activation reveals synergistic potential against Ras pathway dependent cancer cell lines. **(A)** Heatmap of ATARiS gene sensitivity scores of KRAS dependent cell lines (NCI-H358 and MDA-MB-231) and BRAF dependent cell line (A375). Negative values (shaded red) indicate sensitivity of the cell line proliferation to the knockdown of shown genes, whereas positive scores (shaded blue) indicate insensitivity. **(B)** IC_50_ values for various inhibitors in Ras pathway mutant cell lines NCI-H358, MDA-MB231 and A375 grown as serum free 3D spheroids. Compounds were tested either as single agent at a concentration range of 0.2 μM – 10 μM (calmidazolium), 0.6 μM – 40 μM (DT-061), 0.2 μM – 40 μM (trifluoperazine) and 0.6 μM – 40 μM (ARS-1620) or in combination at across the whole concentration range of calmidazolium combined with 1 or 2 μM of DT-061. Data represent mean values ± SD, n = 2. Statistical comparisons were done using the average values across all three cell lines. **(C)** Bliss synergism scores for combinatorial effects of calmidazolium and DT-061 in KRAS- or BRAF-mutant cancer cell lines. Data represent mean values ± SD, n = 2. Positive scores indicate synergistic drug interactions whereas negative scores would denote antagonistic drug interactions. A score of zero would indicate no antagonistic or synergistic effect.

While both calmidazolium or DT-061 alone showed low micromolar inhibition in all cell lines, their combination significantly lowered the average IC_50_ across all tested cell lines (**Figure 1B; Figure S1A-C**). Scoring for synergistic activity of the combination suggested high synergism of these combinations in all three cell lines (**Figure 1C; Figure S1D**). Importantly, a similar level of inhibition as with the combination was achieved using only the PTZ trifluoperazine (**Figure 1B; Figure S1A-C**). Interestingly, both the combination and the single agent trifluoperazine, were as potent as the K-Ras-G12C specific covalent inhibitor ARS-1620 in NCI-H358 cells.

These data, therefore, tentatively suggest that it could be beneficial to employ PTZ derivatives with dual action on CaM and PP2A against cancer cells.

### A covalent phenothiazine derivative binds to CaM and shows enhanced disruption of the cellular K-RasG12V/ CaM interaction

Targeted covalent inhibitors are attracting again more attention as an interesting option in drug development (9). We therefore tested whether a covalent derivative of the PTZ fluphenazine (Flu), with addition of an electrophilic mustard group, fluphenazine mustard (Flu-M), can increase the efficacy of its parental compound.

In order to measure direct inhibition of CaM, we used a fluorescence polarization assay where displacement of a fluorescein-labelled CaM binding peptide, which was derived from plasma membrane Ca^2+^/ATPase (PMCA), by CaM inhibitors is measured (23). Both parental, non-covalently reacting Flu and covalently reacting Flu-M exhibit submicromolar affinities to CaM (**Table 1, Figure 2A**). By contrast, essentially no binding to CaM was detected for the SMAP DT-061, suggesting that it only acts on PP2A.

**Table 1.**
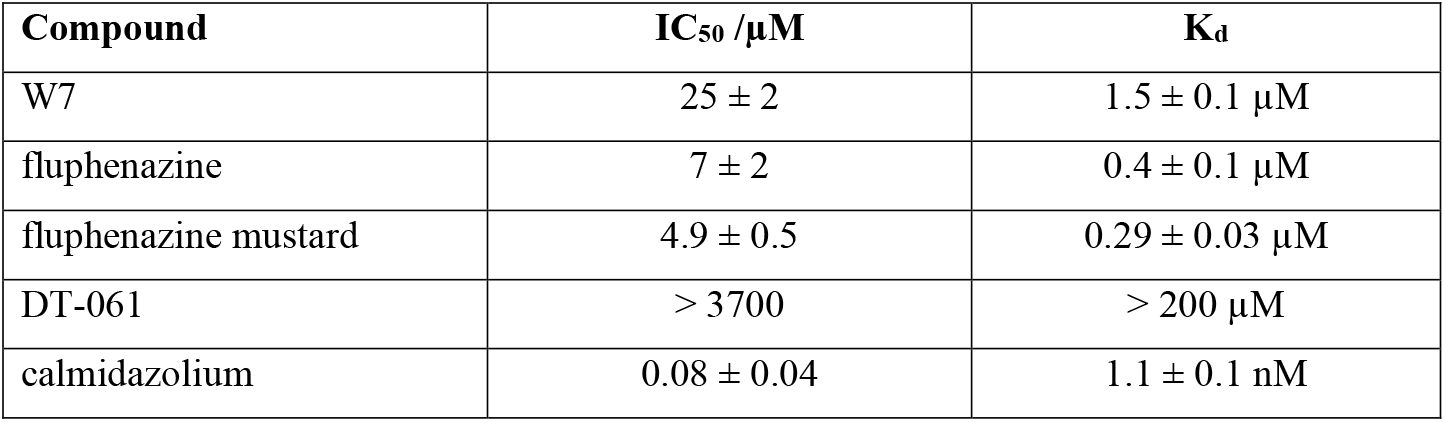
CaM-binding affinity of compounds from fluorescence anisotropy data in Figure 2A (mean ± SD, n = 2).

| Compound | IC<sub>50</sub> /μM | K<sub>d</sub> |
| --- | --- | --- |
| W7 | 25 ± 2 | 1.5 ± 0.1 μM |
| fluphenazine | 7 ± 2 | 0.4 ± 0.1 μM |
| fluphenazine mustard | 4.9 ± 0.5 | 0.29 ± 0.03 μM |
| DT-061 | > 3700 | > 200 μM |
| calmidazolium | 0.08 ± 0.04 | 1.1 ± 0.1 nM |

**Figure 2.**
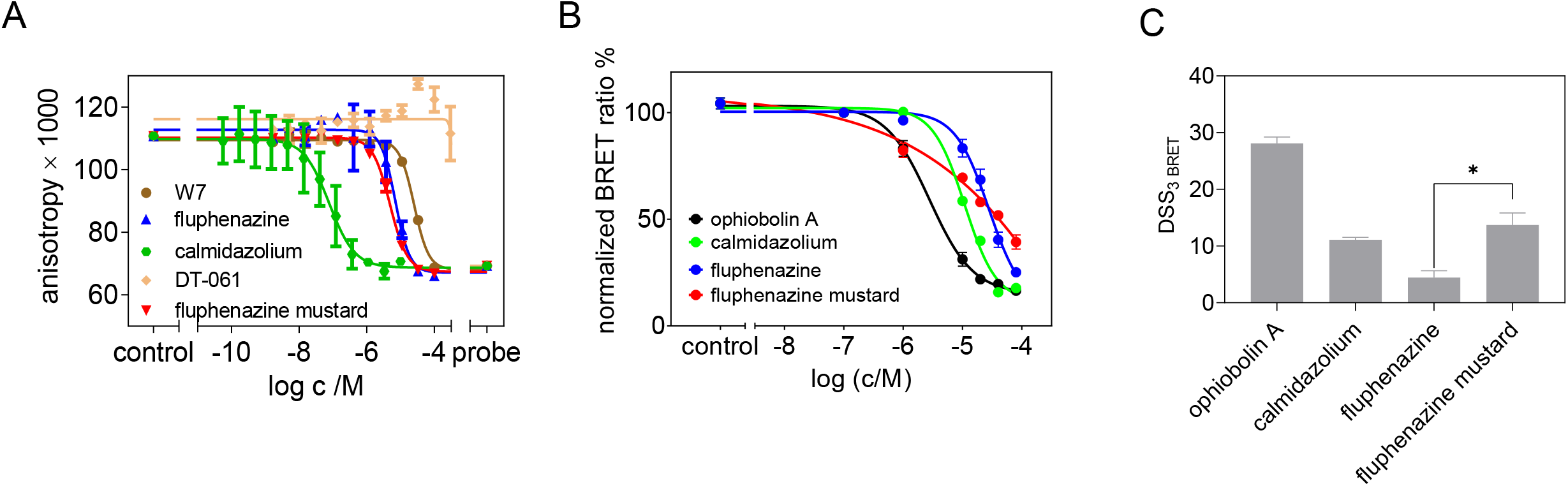
Assessment of the in vitro and in cellulo inhibition of CaM. **(A)** Disruption of a complex of 100 nM CaM and 10 nM F-PMCA peptide by inhibitors was measured in a fluorescence polarization assay. Data represent mean values ± SD, n = 2. **(B)** Dose-response analysis of indicated inhibitors at 0.1 – 80 µM using the Rluc8-K-RasG12V/ GFP2-CaM BRET assay in HEK-293 EBNA cells. The acceptor/ donor plasmid ratio was 9/1. Data represent mean values ± SD, n ≥ 2. **(C)** DSS_3 BRET_ i.e. normalized area under the curve values of dose response data shown in (B).

CaM can act as a trafficking chaperone of K-Ras by shielding the hydrophobic farnesyl tail from the aqueous environment of the cytoplasm, thus effectively increasing the diffusibility of K-Ras (22,40). In order to detect an effect of the CaM inhibitors on the K-Ras/ CaM interaction in cells, we established a BRET assay (22), where we genetically fused Rluc8 to the N-terminus of K-RasG12V and GFP2 to the N-terminus of CaM.

Both the covalently reacting CaM inhibitor OphA and the non-covalent, highly potent calmidazolium reduced the BRET signal when tested across a wider concentration range, in agreement with their CaM inhibiting activity (**Figure 2B**). In order to robustly distinguish the magnitude of these effects, we analyzed the dose response data using a normalized area under the curve measure, the DSS_3 BRET_-score (22,41). These data revealed that addition of the electrophile in Flu-M significantly increased the intracellular activity against CaM/ K-RasG12V as compared to its non-covalent counterpart Flu by more than a factor of two (**Figure 2C**). Yet the potency remained below that of OphA, which however has itself significant off-target activities that may be associated with its broad toxicity spectrum (22).

### Cellular BRET assays confirm Ras membrane organization disrupting properties of fluphenazine mustard

By genetically fusing BRET-donor enabling Rluc8 and the acceptor GFP2 to Ras proteins, we established BRET-assays, which can measure the loss of the functional membrane organization of Ras after inhibition of its trafficking chaperone CaM (22). Tight packing of BRET-luminophore labelled RasG12V in plasma membrane nanoclusters leads to high BRET-levels, similar to what we previously showed (42,43). Loss of nanoclustering, or any process upstream, including proper plasma membrane trafficking or lipid modification of Ras, reduces this nanoclustering-dependent BRET signal. Consistently, treatment with the farnesyl-transferase inhibitor that is known to selectively affect H-Ras more than K-Ras (42), reduced the nanoclustering-dependent BRET signal of H-RasG12V more than that of K-RasG12V (**Figure 3A**,**B**). In agreement, with our previous results (21,22), CaM inhibitors calmidazolium and OphA showed an inverse albeit less pronounced selectivity, reducing the K-RasG12V BRET signal more than that of H-RasG12V (**Figure 3A**,**B**). This was confirmed across a broader concentration range, using the DSS_3 BRET_-score analysis (22) (**Figure 3C; Figure S2A**,**B**).

**Figure 3.**
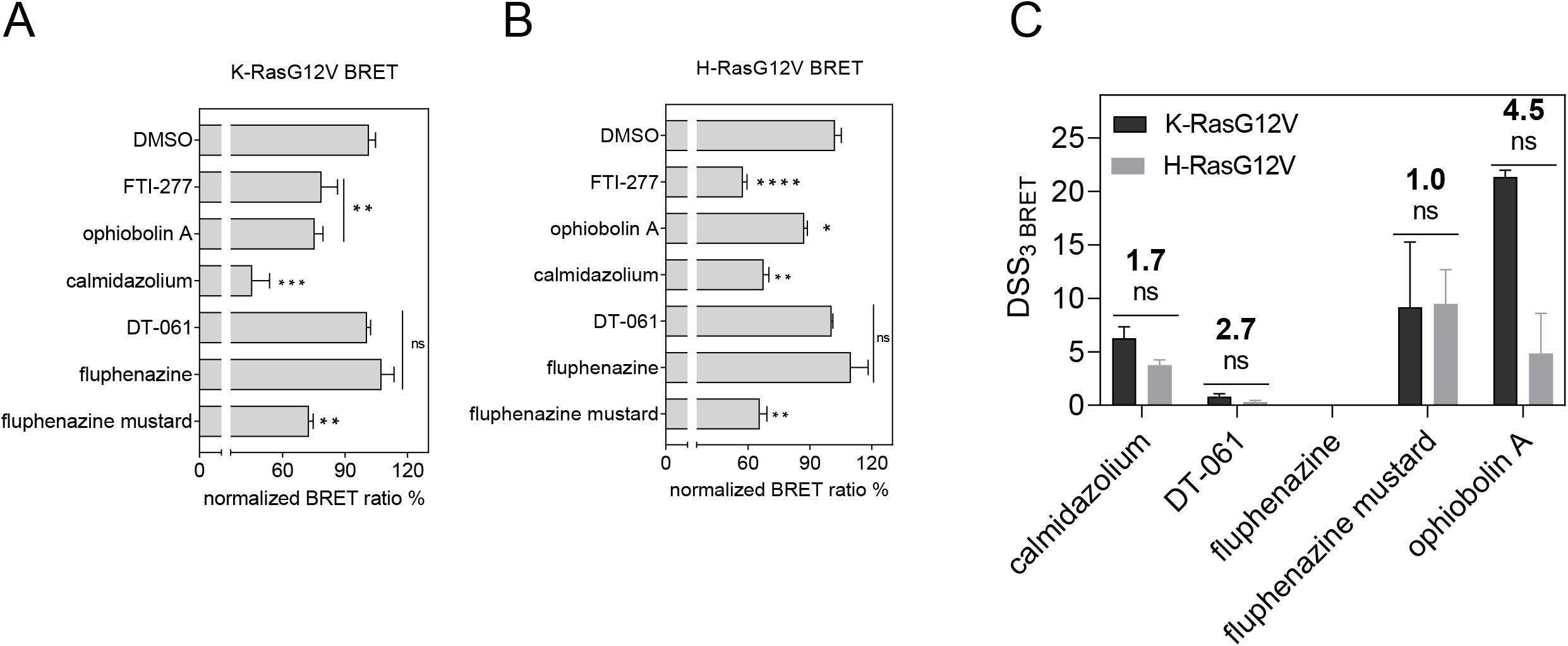
BRET assay reveals increased Ras-potency of fluphenazine mustard in cells. **(A)** Testing of inhibitors at 20 µM for 24 h exposure in KRasG12V **(A)** and H-RasG12V **(B)** nanoclustering-BRET assays. Controls are FTI-277 (1 µM), OphA (2.5 µM), calmidazolium (20 µM) and DT-061 (20 µM). The acceptor/donor plasmid ratio of GFP2- and Rluc8-tagged RasG12V was 4/1. Data represent mean values ± SD, n ≥ 2. **(C)** DSS_3 BRET_ values derived from dose response data for calmidazolium, DT-061, Flu, Flu-M (0.1 μM – 80 µM) and OphA (0.3 μM – 20 µM) using K-RasG12V and H-RasG12V nanoclustering-BRET assays. The acceptor/donor plasmid ratio was 4/1. Data represent mean values ± SD, n ≥ 2.

Neither the PP2A activator DT-061, nor Flu had an effect on the RasG12V-BRET signals (**Figure 3A-C; Figure S2A**,**B**). By contrast, treatment with the covalently-reacting derivative Flu-M led to an even stronger effect than calmidazolium (**Figure 3A-C**), but against both K-RasG12V and H-RasG12V biosensors.

These data suggest that addition of the mustard group can significantly increase the potency of Flu-M against Ras membrane organization in cells.

### ERK-signaling inhibition and strong decrease of cell proliferation by fluphenazine mustard

In agreement with the Ras membrane organization disrupting activity, Flu-M inhibited pERK levels significantly more potently than calmidazolium and DT-061, alone or in combination (**Figure 4A**,**B; Figure S3A**). Hence Flu-M appears to also combine the synergistic activity of the CaM inhibitor and PP2A activator in this context. For comparison, MAPK-signaling inhibition by the clinically employed KRAS-G12C inhibitor AMG-510 or MEK inhibitor trametinib was assessed (**Figure 4A**,**B; Figure S3A**). Both of these inhibitors were at least 20-times more potent than Flu-M in this assay.

**Figure 4.**
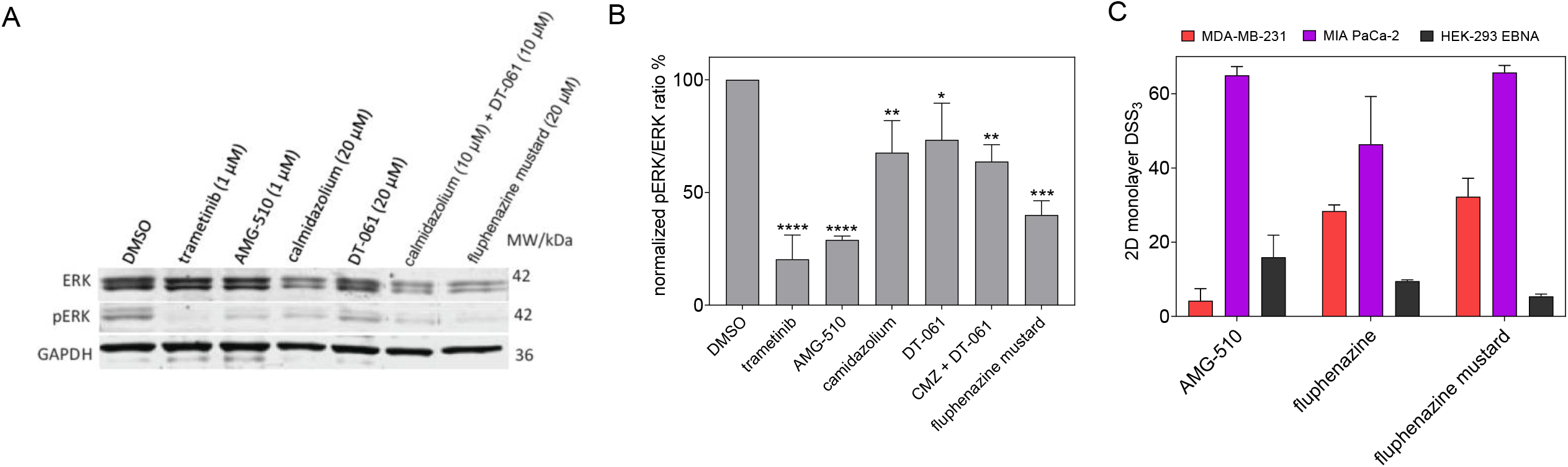
Signaling and anti-proliferative effects. **(A)** MAPK signaling output was measured in MIA PaCa-2 cells after inhibitor treatment and EGF stimulation. DMSO is 0.2 % (v/v) vehicle control. Representative blot from three independent biological repeats. **(B)** Densitometric quantification of the pERK/ERK ratio of all three biological repeats (mean values ± SD) **(C)** DSS_3_ measuring the anti-proliferative effects of AMG-510 (0.003 – 40 μM), fluphenazine (0.6 – 80 μM) and fluphenazine mustard (0.6 – 80 μM). Results represent mean values ± SD, n = 3.

However, when tested for its effect on cell proliferation, both Flu and Flu-M appeared relatively potent as compared to AMG-510, with Flu-M being significantly more potent against MIA PaCa-2 cell proliferation than Flu (**Figure 4C; Figure S3B-D**). Given that Flu-M has a moderate inhibitory activity on pERK-levels, the relatively potent inhibition of cell proliferation is in agreement with multiple targeting mechanisms of this compound.

## Discussion

Here we provided evidence that PTZ can integrate the synergistic activities of CaM inhibition and PP2A activation within one compound. This activity can be potentiated using a PTZ, such as Flu-M, equipped with an electrophile that can covalently inactivate one or both of these targets. At this point, it is not clear, whether the covalent derivative Flu-M also covalently engages PP2A. CaM inhibition by PTZ is supported by in vitro and in cellulo data. By contrast, the SMAP DT-061 does not inhibit CaM. Given that CaM is a trafficking chaperone in particular of K-Ras, CaM inhibition disrupts functional Ras-membrane organization and downstream MAPK-output. This is supported by the relatively high potency against 2D proliferation of MIA PaCa-2 cells, while it is also in line with the engagement of several targets, such as PP2A.

Importantly, addition of the electrophile did not increase unspecific toxicity in cells, as non-transformed HEK293 cells were > 2-fold less affected by Flu-M than by Flu (**Figure 4C**). Taken together with the more potent inhibition of strongly KRAS-dependent MIA PaCa-2 cells and the increased CaM inhibition in cells (**Figures 2C**), a significant contribution of CaM inhibition in the cell killing phenotype is apparent. Furthermore, the dual activity of Flu-M against K-RasG12V and H-RasG12V membrane anchorage BRET may also explain the high potency (**Figure 3C**). It is currently not clear, why also the H-RasG12V BRET-signal was affected, while we previously noted a clear K-Ras selectivity of non-covalent and covalent CaM inhibitors (22). It may suggest that other targets are engaged by this compound that affect membrane organization of both Ras isoforms.

While our study is quite limited, it suggests that preservation of the CaM inhibitory activity of PTZ derivatives that can also activate PP2A is desirable. Our data show however, that DT-061 does not bind to CaM and would therefore have lost the synergistic potential. We propose that the preservation of the investigated polypharmacology of PTZ could allow for a higher, yet selective cancer cell killing activity.

This is supported by substantial evidence for the anti-cancer activity of PTZ. Several studies have shown that PTZs inhibit the growth of various cancer types including glioblastomas, melanoma, colorectal cancer and acute lymphoblastic leukemia (38,44,45). The ability of PTZs to effect multiple biological consequences such as the disruption of membrane functions, DNA repair, cell cycle regulation, apoptosis and multiple signaling pathways indicates the potential of this class of compounds as viable anti-cancer agents (46,47). Interestingly, the dopamine receptor mRNA and protein expression levels are elevated in a variety of cancers including cervical, esophageal, glioma, breast cancer and lung cancer (48-52). Therefore, also this target may play a role in the anti-cancer effect of PTZ.

We could show that the mustard derivative of the PTZ fluphenazine, Flu-M, which has a quite reactive electrophile, affords a better cancer cell selectivity than its non-covalent counterpart. It can be expected that further optimizations here, could allow for improved chemical selectivity, for instance by using a softer electrophile such as an acrylamide. One particular advantage of covalent inhibitors is that the covalent bonding step might actually improve selectivity for a desired target, as only that target may exhibit proper nucleophile constellation near the non-covalent initial binding site.

Alternative PP2A activators have been identified, such as the sphingosine 1-phosphate receptor modulator FTY720 (Fingolimod), which is FDA approved for multiple-sclerosis. It binds to the endogenous PP2A inhibitor SET, thus efficaciously reducing leukemic burden in mice (53). This potential has been recapitulated by an improved derivative, CM-1231, which shows no more cardiotoxicity as compared to Fingolimod and could be developed for treatment of leukemic patients with SET overexpression (54). It remains to be seen whether Fingolimod derivatives or PTZ derivatives can be developed into better PP2A activators.

Based on our data, we suggest to develop PTZ derivatives that still bind to CaM and activate PP2A, while being devoid of any neurological side-effect. Such compounds would combine the synergistic activity of CaM inhibition and PP2A activation in one molecule and could thus possess an increased and selective cancer cell killing activity.

## Experimental procedures

### Reagents

Sources of the compounds used in the study are given in parentheses, next to compounds name. Fluphenazine (Sigma, F0280000); Fluphenazine mustard (Enzo, BML-CA325-0050); Trifluoperazine (Sigma, 1686003); DT-061 (MedChem Express, HY-112929); Ophiobolin A (Santa Cruz, sc-202266); Calmidazolium (Santa Cruz, sc-201494); FTI-277 (BioVision, 2874); AMG-510 (MedChem Express, HY-114277); trametinib (Bio-connect, SC-364639). DMSO was from PanReac-AppliChem (cat. no. A3672, ITW Reagents).

### ATARiS gene dependence score

ATARiS gene dependency scores for each gene of interest were extracted from the publicly available database of the project DRIVE (https://oncologynibr.shinyapps.io/drive/) (55). The normalized viability data for the siRNA knockdown of each gene of interest were downloaded from the project DRIVE and a double gradient heatmap plot was generated using Prism (GraphPad).

### 3D Spheroid Assays

3D spheroid formation assays were performed in 96-well low-attachment, suspension culture plates (cat. no. 655185, Cellstar, Greiner Bio-One) under serum free condition. About 1000 cells per well were seeded in 50 µL RPMI medium (cat. no. 52400-025, Gibco, ThermoFisher Scientific) containing 0.5 % (v/v) MethoCult (cat. no. SF H4636, Stemcell technologies), 1X B27 (cat. no. 17504044, Gibco, ThermoFisher Scientific), 25 ng/mL EGF (cat. no. E9644, Sigma-Aldrich) and 25 ng/ mL FGF (cat. no. RP-8628, ThermoFisher Scientific). Cells were cultured for 3 days and then treated with compounds or vehicle control (DMSO 0.1 % (v/v) in growth medium) for another 3 days. The cells were supplemented with fresh growth medium on the third day together with the drug treatment. Spheroid formation efficiency was analyzed by alamarBlue assay reagent (cat. no. DAL1100, ThermoFisher Scientific).

A 10 % (v/v) final volume of alamarBlue reagent was added to each well of the plate and incubated for 4 h at 37 °C. Then the fluorescence intensity (λ_excitation_ 560 ± 5 nm and λ_emission_ 590 ± 5 nm) was measured using the FLUOstar OPTIMA plate reader (BMG Labtech, Germany). The obtained fluorescence intensity data were normalized to vehicle control corresponding to 100 % sphere formation and the signal after 100 μM benzethonium chloride (cat. no. 53751, Sigma-Aldrich) treatment, which killed all cells (i.e. maximum inhibition of sphere formation).

### Bliss synergism experiments

To assess the synergistic potential of our compounds, we performed a full dose response analysis of one inhibitor (calmidazolium) and maintained a fixed concentration of the other inhibitor (DT-061) at either 1 or 2 µM. The drug response profiles obtained for the combinations were then compared against the profiles of each single agent using the SynergyFinder platform (56). SynergyFinder (https://synergyfinder.fimm.fi) is a stand-alone web-application for interactive analysis and visualization of drug combination screening data (56). We employed the Bliss model (57), which generates the multiplicative effects of single agents as if they acted independently in scoring our pairwise combinations

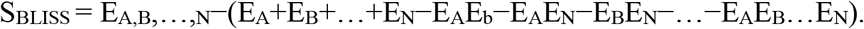

Here, E_A_,E_B_,…,E_N_ are the measured responses of the single drugs, while *b* is the doses of the single drugs required to produce the combination effect E_A_,_B_,…,_N_.

### Drug sensitivity score (DSS) analysis

The Drug Sensitivity Score (DSS) is a more robust parameter than IC_50_ or EC_50_ and measures essentially the normalized area under the curve of dose-response data (41). The normalized % inhibition data of the BRET assay or the raw intensity data of the 2D-monolayer assay were uploaded to the DSS pipeline website, Breeze (https://breeze.fimm.fi/) (22,58). The output file from the Breeze platform containing DSS_3_ and several other drug sensitivity measures including IC_50_ or EC_50_ and AUC was downloaded after the simulation.

### 2D cell viability assays

HEK-293 EBNA, MDA-MB-23 and MIA PaCa-2 cells, cultured in complete DMEM and RPMI medium i.e supplemented with 10 % (v/v) FBS (cat. no. 10270-098, Gibco, ThermoFisher Scientific), 2 mM L-glutamine (cat. no. 25030-024, ThermoFisher Scientific), respectively, were plated into 96-well F-bottom cell culture plates (cat. no. 655180, Cellstar, Greiner Bio-One) at a density of 1000 cells per well and grown for 24 h. Compounds at indicated concentration or DMSO at 0.2 % (v/v) in growth medium were treated for 72 h then the cell viability was measured using alamarBlue assay. A 10 % (v/v) final volume of alamarBlue reagent was added to each well of the plate and incubated for 4 h at 37 °C. Then the fluorescence intensity (λ_excitation_ 560 ± 5 nm and λ_emission_ 590 ± 5 nm) was measured using the FLUOstar OPTIMA plate reader (BMG Labtech). The obtained fluorescence intensity data were normalized to vehicle control (100 % viability).

### Fluorescence polarization assay

The fluorescence polarization assay was performed as previously described (22,23). A complex of recombinant bovine CaM (cat. no. 208690, Merck), with an amino acid sequence identical to the human isoform, and fluorescein-labelled PMCA peptide was added to a 3-fold dilution series of inhibitor in an assay buffer (20 mM Tris Cl pH 7.5, 50 mM NaCl, 1 mM CaCl_2_ and 0.005 % (v/v) Tween 20) in a black low volume round bottom 384-well plate (cat. no. 4514, Corning). After 24 h incubation, the fluorescence anisotropy (λ_excitation_ 482 ± 8 nm and λ_emission_ 530 ± 20 nm) was recorded in a Clariostar (BMG labtech) plate reader. The IC_50_ value was derived by fitting the log concentration of inhibitor vs fluorescence anisotropy signal in Prism (GraphPad) software. The IC_50_ value was converted into *K*_d_ as described previously (59).

### BRET assays

The detailed BRET methodology and plasmids encoding Rluc8-K-RasG12V, Rluc8-H-RasG12V, GFP2-K-RasG12V, GFP2-H-RasG12V and GFP2-CaM were recently described (22). About 100,000 to 150,000 HEK-293 EBNA (60) cells were seeded per well of a 12-well plate in 1 mL of DMEM containing 10 % (v/v) FBS and 0.5 mM L-glutamine and grown overnight. Next day 1 µg of BRET sensor plasmids were transfected using 3 µL of jetPRIME reagent (cat. no. 114-75, Polyplus). The ratio of acceptor to donor plasmid (A/D plasmid ratio) for each experiment is indicated in the figure legends. After 24 h of transfection, cells were treated with vehicle control (DMSO 0.2 % (v/v) in growth medium) or compounds at the specified concentration for 24 h. Then the cells were collected, washed and re-plated in PBS (cat. no. 14190-094, Gibco, ThermoFisher Scientific) into flat bottom, white 96-well plates (cat. no. 236108, Nunc, ThermoFisher Scientific). First the fluorescence intensity (λ_excitation_ 405 ± 10 nm and λ_emission_ 515 ± 10 nm) of GFP2 was measured as it is directly proportional to the acceptor concentration. Next the BRET readings were taken in well-mode by adding coelenterazine 400a (cat. no. C-320, GoldBio) to a final concentration of 10 µM and luminescence emission intensities were simultaneously recorded at 410 ± 40 nm and at 515 ± 15 nm. The raw BRET ratio was calculated as the BRET signal measured at 515 nm divided by the emission signal measured at 410 nm. The BRET ratio was obtained by subtracting the raw BRET ratio from the background BRET signal measured for cells expressing only the donor.

### Western blotting

MIA PaCa-2 cells were treated with various inhibitors for 2 h and then stimulated for 10 min with EGF (50 ng/ mL), lysed with cell lysis buffer (50 mM Tris, 150 mM NaCl, 0.1 % (v/v) SDS, 1 % (v/v) Triton X-100, 1 % (v/v) NP40 and 1 x cOmplete Protease inhibitor cocktail (Roche)) by incubating on ice for 30 min. The clarified cell lysates were mixed with Laemmli buffer containing 0.1 % (v/v) bromophenol blue heated for 5 min at 95 °C and resolved on a 12 % home-made acrylamide gel. Proteins were subsequently transferred to a nitrocellulose membrane (BIO-RAD) using the Trans-Blot Turbo Transfer system (BIO-RAD). The blotted membranes were incubated for 1 h in blocking buffer (5 % (w/v) BSA in Tris-buffered saline (TBS)–T (Tween 0.1 % (v/v))) and treated with primary antibodies for pERK 1:400 (Santa cruz, sc-81503), ERK 1:1000 (Cell Signaling, 9102). A mouse antibody against GAPDH 1:2000 (Sigma, G8795) was used as the loading control. After overnight incubation, the membrane was washed 3 × 5 min in TBS–T buffer the following day and then treated with the corresponding anti-rabbit and anti-mouse secondary antibodies (1:10000 dilution). The proteins were detected using a LI-COR ODYSSEY CLx system (Westburg). The level of proteins was densitometrically quantified from western blot membranes using ImageLab software (BIO-RAD).

### Data analysis

All data analysis was performed using Prism (GraphPad) version 9. The number of independent biological repeats, n, for each data set is provided in the relevant figure legend. Unless otherwise stated, statistical significance was evaluated using one-way ANOVA. A p-value of < 0.05 is considered statistically significant and the statistical significance levels are annotated as: * = p < 0.05; ** = p < 0.01; *** = p < 0.001; **** = p < 0.0001, or ns = not significant.

## Supporting Information

This article contains supporting information, available online.

## Acknowledgements

We thank Ms. Marie Catillon for technical support and Dr. Karolina Pavic for critical comments on the manuscript.

## Conflict of interest

The authors declare that they have no conflicts of interests.

## References

1. Jaszczyszyn, A., Gąsiorowski, K., Świątek, P., Malinka, W., Cieślik-Boczula, K., Petrus, J., and Czarnik-Matusewicz, B. (2012) Chemical structure of phenothiazines and their biological activity. Pharmacol Rep 64, 16–23

2. Prozialeck, W. C., and Weiss, B. (1982) Inhibition of calmodulin by phenothiazines and related drugs: structure-activity relationships. J. Pharmacol. Exp. Ther. 222, 509–516

3. Berchtold, M. W., and Villalobo, A. (2014) The many faces of calmodulin in cell proliferation, programmed cell death, autophagy, and cancer. Biochim Biophys Acta 1843, 398–435

4. Sharma, R. K., and Parameswaran, S. (2018) Calmodulin-binding proteins: A journey of 40 years. Cell Calcium 75, 89–100

5. Hait, W. N., and Lazo, J. S. (1986) Calmodulin: a potential target for cancer chemotherapeutic agents. J Clin Oncol 4, 994–1012

6. Hait, W. N., Byrne, T. N., Piepmeier, J., Durivage, H. J., Choudhury, S., Davis, C. A., and Gates, J. A. (1990) The effect of calmodulin inhibitors with bleomycin on the treatment of patients with high grade gliomas. Cancer Res 50, 6636–6640

7. Kong Au, T., and Chow Leung, P. (1998) Identification of the binding and inhibition sites in the calmodulin molecule for ophiobolin A by site-directed mutagenesis. Plant Physiol 118, 965–973

8. Faust, F. M., Slisz, M., and Jarrett, H. W. (1987) Calmodulin is labeled at lysine 148 by a chemically reactive phenothiazine. J Biol Chem 262, 1938–1941

9. Singh, J., Petter, R. C., Baillie, T. A., and Whitty, A. (2011) The resurgence of covalent drugs. Nat Rev Drug Discov 10, 307–317

10. Hait, W. N., Glazer, L., Kaiser, C., Cross, J., and Kennedy, K. A. (1987) Pharmacological properties of fluphenazine-mustard, an irreversible calmodulin antagonist. Mol. Pharmacol. 32, 404–409

11. Singh, R. K., Kumar, S., Prasad, D. N., and Bhardwaj, T. R. (2018) Therapeutic journery of nitrogen mustard as alkylating anticancer agents: Historic to future perspectives. Eur J Med Chem 151, 401–433

12. Ostrem, J. M., Peters, U., Sos, M. L., Wells, J. A., and Shokat, K. M. (2013) K-Ras(G12C) inhibitors allosterically control GTP affinity and effector interactions. Nature 503, 548–551

13. Janes, M. R., Zhang, J., Li, L. S., Hansen, R., Peters, U., Guo, X., Chen, Y., Babbar, A., Firdaus, S. J., Darjania, L., Feng, J., Chen, J. H., Li, S., Li, S., Long, Y. O., Thach, C., Liu, Y., Zarieh, A., Ely, T., Kucharski, J. M., Kessler, L. V., Wu, T., Yu, K., Wang, Y., Yao, Y., Deng, X., Zarrinkar, P. P., Brehmer, D., Dhanak, D., Lorenzi, M. V., Hu-Lowe, D., Patricelli, M. P., Ren, P., and Liu, Y. (2018) Targeting KRAS Mutant Cancers with a Covalent G12C-Specific Inhibitor. Cell 172, 578–589 e517

14. Canon, J., Rex, K., Saiki, A. Y., Mohr, C., Cooke, K., Bagal, D., Gaida, K., Holt, T., Knutson, C. G., Koppada, N., Lanman, B. A., Werner, J., Rapaport, A. S., San Miguel, T., Ortiz, R., Osgood, T., Sun, J. R., Zhu, X., McCarter, J. D., Volak, L. P., Houk, B. E., Fakih, M. G., O’Neil, B. H., Price, T. J., Falchook, G. S., Desai, J., Kuo, J., Govindan, R., Hong, D. S., Ouyang, W., Henary, H., Arvedson, T., Cee, V. J., and Lipford, J. R. (2019) The clinical KRAS(G12C) inhibitor AMG 510 drives anti-tumour immunity. Nature 575, 217–223

15. Hallin, J., Engstrom, L. D., Hargis, L., Calinisan, A., Aranda, R., Briere, D. M., Sudhakar, N., Bowcut, V., Baer, B. R., Ballard, J. A., Burkard, M. R., Fell, J. B., Fischer, J. P., Vigers, G. P., Xue, Y., Gatto, S., Fernandez-Banet, J., Pavlicek, A., Velastagui, K., Chao, R. C., Barton, J., Pierobon, M., Baldelli, E., Patricoin, E. F., 3rd, Cassidy, D.P., Marx, M. A., Rybkin, II, Johnson, M. L., Ou, S. I., Lito, P., Papadopoulos, K. P., Janne, P. A., Olson, P., and Christensen, J. G. (2020) The KRAS(G12C) Inhibitor MRTX849 Provides Insight toward Therapeutic Susceptibility of KRAS-Mutant Cancers in Mouse Models and Patients. Cancer Discovery 10, 54–71

16. Hong, D. S., Fakih, M. G., Strickler, J. H., Desai, J., Durm, G. A., Shapiro, G. I., Falchook, G. S., Price, T. J., Sacher, A., Denlinger, C. S., Bang, Y. J., Dy, G. K., Krauss, J. C., Kuboki, Y., Kuo, J. C., Coveler, A. L., Park, K., Kim, T. W., Barlesi, F., Munster, P. N., Ramalingam, S. S., Burns, T. F., Meric-Bernstam, F., Henary, H., Ngang, J., Ngarmchamnanrith, G., Kim, J., Houk, B. E., Canon, J., Lipford, J. R., Friberg, G., Lito, P., Govindan, R., and Li, B. T. (2020) KRAS(G12C) Inhibition with Sotorasib in Advanced Solid Tumors. N Engl J Med 383, 1207–1217

17. Spiegel, J., Cromm, P. M., Zimmermann, G., Grossmann, T. N., and Waldmann, H. (2014) Small-molecule modulation of Ras signaling. Nat Chem Biol 10, 613–622

18. Grant, B. M. M., Enomoto, M., Ikura, M., and Marshall, C. B. (2020) A Non-Canonical Calmodulin Target Motif Comprising a Polybasic Region and Lipidated Terminal Residue Regulates Localization. IJMS 21

19. Chippalkatti, R., and Abankwa, D. (2021) Promotion of cancer cell stemness by Ras. Biochem Soc Trans 49, 467–476

20. Wang, M. T., Holderfield, M., Galeas, J., Delrosario, R., To, M. D., Balmain, A., and McCormick, F. (2015) K-Ras Promotes Tumorigenicity through Suppression of Non-canonical Wnt Signaling. Cell 163, 1237–1251

21. Najumudeen, A. K., Jaiswal, A., Lectez, B., Oetken-Lindholm, C., Guzman, C., Siljamaki, E., Posada, I. M., Lacey, E., Aittokallio, T., and Abankwa, D. (2016) Cancer stem cell drugs target K-ras signaling in a stemness context. Oncogene 35, 5248–5262

22. Okutachi, S., Manoharan, G. B., Kiriazis, A., Laurini, C., Catillon, M., McCormick, F., Yli-Kauhaluoma, J., and Abankwa, D. (2021) A covalent calmodulin inhibitor as a tool to study cellular mechanisms of K-Ras-driven stemness. Frontiers in Cell and Developmental Biology; DOI: 10.3389/fcell.2021.665673

23. Manoharan, G. B., Kopra, K., Eskonen, V., Härmä, H., and Abankwa, D. (2019) High-throughput amenable fluorescence-assays to screen for calmodulin-inhibitors. Anal Biochem 572, 25–32

24. Perrotti, D., and Neviani, P. (2013) Protein phosphatase 2A: a target for anticancer therapy. Lancet Oncol 14, e229–238

25. Sangodkar, J., Farrington, C. C., McClinch, K., Galsky, M. D., Kastrinsky, D. B., and Narla, G. (2016) All roads lead to PP2A: exploiting the therapeutic potential of this phosphatase. FEBS J 283, 1004–1024

26. Rangarajan, A., Hong, S. J., Gifford, A., and Weinberg, R. A. (2004) Species- and cell type-specific requirements for cellular transformation. Cancer Cell 6, 171–183

27. Shi, Y. (2009) Serine/threonine phosphatases: mechanism through structure. Cell 139, 468–484

28. Fowle, H., Zhao, Z., and Grana, X. (2019) PP2A holoenzymes, substrate specificity driving cellular functions and deregulation in cancer. Adv Cancer Res 144, 55–93

29. Jackson, J. B., and Pallas, D. C. (2012) Circumventing cellular control of PP2A by methylation promotes transformation in an Akt-dependent manner. Neoplasia 14, 585–599

30. Junttila, M. R., Puustinen, P., Niemela, M., Ahola, R., Arnold, H., Bottzauw, T., Ala-aho, R., Nielsen, C., Ivaska, J., Taya, Y., Lu, S. L., Lin, S., Chan, E. K., Wang, X. J., Grenman, R., Kast, J., Kallunki, T., Sears, R., Kahari, V. M., and Westermarck, J. (2007) CIP2A inhibits PP2A in human malignancies. Cell 130, 51–62

31. Wang, J., Okkeri, J., Pavic, K., Wang, Z., Kauko, O., Halonen, T., Sarek, G., Ojala, P. M., Rao, Z., Xu, W., and Westermarck, J. (2017) Oncoprotein CIP2A is stabilized via interaction with tumor suppressor PP2A/B56. EMBO Rep 18, 437–450

32. Zhou, X., Updegraff, B. L., Guo, Y., Peyton, M., Girard, L., Larsen, J. E., Xie, X. J., Zhou, Y., Hwang, T. H., Xie, Y., Rodriguez-Canales, J., Villalobos, P., Behrens, C., Wistuba, II, Minna, J. D., and O’Donnell, K. A. (2017) PROTOCADHERIN 7 Acts through SET and PP2A to Potentiate MAPK Signaling by EGFR and KRAS during Lung Tumorigenesis. Cancer Res 77, 187–197

33. Kauko, O., O’Connor, C. M., Kulesskiy, E., Sangodkar, J., Aakula, A., Izadmehr, S., Yetukuri, L., Yadav, B., Padzik, A., Laajala, T. D., Haapaniemi, P., Momeny, M., Varila, T., Ohlmeyer, M., Aittokallio, T., Wennerberg, K., Narla, G., and Westermarck, J. (2018) PP2A inhibition is a druggable MEK inhibitor resistance mechanism in KRAS-mutant lung cancer cells. Sci Transl Med 10

34. Sangodkar, J., Perl, A., Tohme, R., Kiselar, J., Kastrinsky, D. B., Zaware, N., Izadmehr, S., Mazhar, S., Wiredja, D. D., O’Connor, C. M., Hoon, D., Dhawan, N. S., Schlatzer, D., Yao, S., Leonard, D., Borczuk, A. C., Gokulrangan, G., Wang, L., Svenson, E., Farrington, C. C., Yuan, E., Avelar, R. A., Stachnik, A., Smith, B., Gidwani, V., Giannini, H. M., McQuaid, D., McClinch, K., Wang, Z., Levine, A. C., Sears, R. C., Chen, E. Y., Duan, Q., Datt, M., Haider, S., Ma’ayan, A., DiFeo, A., Sharma, N., Galsky, M. D., Brautigan, D. L., Ioannou, Y. A., Xu, W., Chance, M. R., Ohlmeyer, M., and Narla, G. (2017) Activation of tumor suppressor protein PP2A inhibits KRAS-driven tumor growth. J Clin Invest 127, 2081–2090

35. Kastrinsky, D. B., Sangodkar, J., Zaware, N., Izadmehr, S., Dhawan, N. S., Narla, G., and Ohlmeyer, M. (2015) Reengineered tricyclic anti-cancer agents. Bioorg Med Chem 23, 6528–6534

36. Leonard, D., Huang, W., Izadmehr, S., O’Connor, C. M., Wiredja, D. D., Wang, Z., Zaware, N., Chen, Y., Schlatzer, D. M., Kiselar, J., Vasireddi, N., Schuchner, S., Perl, A. L., Galsky, M. D., Xu, W., Brautigan, D. L., Ogris, E., Taylor, D. J., and Narla, G. (2020) Selective PP2A Enhancement through Biased Heterotrimer Stabilization. Cell 181, 688–701 e616

37. Morita, K., He, S., Nowak, R. P., Wang, J., Zimmerman, M. W., Fu, C., Durbin, A. D., Martel, M. W., Prutsch, N., Gray, N. S., Fischer, E. S., and Look, A. T. (2020) Allosteric Activators of Protein Phosphatase 2A Display Broad Antitumor Activity Mediated by Dephosphorylation of MYBL2. Cell 181, 702–715 e720

38. Wu, C. H., Bai, L. Y., Tsai, M. H., Chu, P. C., Chiu, C. F., Chen, M. Y., Chiu, S. J., Chiang, J. H., and Weng, J. R. (2016) Pharmacological exploitation of the phenothiazine antipsychotics to develop novel antitumor agents-A drug repurposing strategy. Sci Rep 6, 27540

39. Motohashi, N., Kawase, M., Saito, S., and Sakagami, H. (2000) Antitumor potential and possible targets of phenothiazine-related compounds. Curr Drug Targets 1, 237–245

40. Grant, B. M. M., Enomoto, M., Back, S. I., Lee, K. Y., Gebregiworgis, T., Ishiyama, N., Ikura, M., and Marshall, C. B. (2020) Calmodulin disrupts plasma membrane localization of farnesylated KRAS4b by sequestering its lipid moiety. Science Signaling 13, eaaz0344

41. Yadav, B., Pemovska, T., Szwajda, A., Kulesskiy, E., Kontro, M., Karjalainen, R., Majumder, M. M., Malani, D., Murumagi, A., Knowles, J., Porkka, K., Heckman, C., Kallioniemi, O., Wennerberg, K., and Aittokallio, T. (2014) Quantitative scoring of differential drug sensitivity for individually optimized anticancer therapies. Sci Rep 4, 5193

42. Kohnke, M., Schmitt, S., Ariotti, N., Piggott, A. M., Parton, R. G., Lacey, E., Capon, R. J., Alexandrov, K., and Abankwa, D. (2012) Design and application of in vivo FRET biosensors to identify protein prenylation and nanoclustering inhibitors. Chemistry & Biology 19, 866–874

43. Najumudeen, A. K., Kohnke, M., Solman, M., Alexandrov, K., and Abankwa, D. (2013) Cellular FRET-Biosensors to Detect Membrane Targeting Inhibitors of N-Myristoylated Proteins. PLoS One 8, e66425

44. Gutierrez, A., Pan, L., Groen, R. W., Baleydier, F., Kentsis, A., Marineau, J., Grebliunaite, R., Kozakewich, E., Reed, C., Pflumio, F., Poglio, S., Uzan, B., Clemons, P., VerPlank, L., An, F., Burbank, J., Norton, S., Tolliday, N., Steen, H., Weng, A. P., Yuan, H., Bradner, J. E., Mitsiades, C., Look, A. T., and Aster, J. C. (2014) Phenothiazines induce PP2A-mediated apoptosis in T cell acute lymphoblastic leukemia. J Clin Invest 124, 644–655

45. Jiang, X., Chen, Z., Shen, G., Jiang, Y., Wu, L., Li, X., Wang, G., and Yin, T. (2018) Psychotropic agent thioridazine elicits potent in vitro and in vivo anti-melanoma effects. Biomed Pharmacother 97, 833–837

46. Sudeshna, G., and Parimal, K. (2010) Multiple non-psychiatric effects of phenothiazines: a review. Eur J Pharmacol 648, 6–14

47. Motohashi, N., Kawase, M., Satoh, K., and Sakagami, H. (2006) Cytotoxic potential of phenothiazines. Curr Drug Targets 7, 1055–1066

48. Li, L., Miyamoto, M., Ebihara, Y., Mega, S., Takahashi, R., Hase, R., Kaneko, H., Kadoya, M., Itoh, T., Shichinohe, T., Hirano, S., and Kondo, S. (2006) DRD2/DARPP-32 expression correlates with lymph node metastasis and tumor progression in patients with esophageal squamous cell carcinoma. World J Surg 30, 1672-1679; discussion 1680-1671

49. Li, J., Zhu, S., Kozono, D., Ng, K., Futalan, D., Shen, Y., Akers, J. C., Steed, T., Kushwaha, D., Schlabach, M., Carter, B. S., Kwon, C. H., Furnari, F., Cavenee, W., Elledge, S., and Chen, C. C. (2014) Genome-wide shRNA screen revealed integrated mitogenic signaling between dopamine receptor D2 (DRD2) and epidermal growth factor receptor (EGFR) in glioblastoma. Oncotarget 5, 882–893

50. Hoeppner, L. H., Wang, Y., Sharma, A., Javeed, N., Van Keulen, V. P., Wang, E., Yang, P., Roden, A. C., Peikert, T., Molina, J. R., and Mukhopadhyay, D. (2015) Dopamine D2 receptor agonists inhibit lung cancer progression by reducing angiogenesis and tumor infiltrating myeloid derived suppressor cells. Mol Oncol 9, 270–281

51. Pornour, M., Ahangari, G., Hejazi, S. H., Ahmadkhaniha, H. R., and Akbari, M. E. (2014) Dopamine receptor gene (DRD1-DRD5) expression changes as stress factors associated with breast cancer. Asian Pac J Cancer Prev 15, 10339–10343

52. Weissenrieder, J. S., Neighbors, J. D., Mailman, R. B., and Hohl, R. J. (2019) Cancer and the Dopamine D2 Receptor: A Pharmacological Perspective. J Pharmacol Exp Ther 370, 111–126

53. Oaks, J. J., Santhanam, R., Walker, C. J., Roof, S., Harb, J. G., Ferenchak, G., Eisfeld, A. K., Van Brocklyn, J. R., Briesewitz, R., Saddoughi, S. A., Nagata, K., Bittman, R., Caligiuri, M. A., Abdel-Wahab, O., Levine, R., Arlinghaus, R. B., Quintas-Cardama, A., Goldman, J. M., Apperley, J., Reid, A., Milojkovic, D., Ziolo, M. T., Marcucci, G., Ogretmen, B., Neviani, P., and Perrotti, D. (2013) Antagonistic activities of the immunomodulator and PP2A-activating drug FTY720 (Fingolimod, Gilenya) in Jak2-driven hematologic malignancies. Blood 122, 1923–1934

54. Vicente, C., Arriazu, E., Martinez-Balsalobre, E., Peris, I., Marcotegui, N., Garcia-Ramirez, P., Pippa, R., Rabal, O., Oyarzabal, J., Guruceaga, E., Prosper, F., Mateos, M. C., Cayuela, M. L., and Odero, M. D. (2020) A novel FTY720 analogue targets SET-PP2A interaction and inhibits growth of acute myeloid leukemia cells without inducing cardiac toxicity. Cancer Lett 468, 1–13

55. McDonald, E. R., 3rd, de Weck, A., Schlabach, M. R., Billy, E., Mavrakis, K. J., Hoffman, G. R., Belur, D., Castelletti, D., Frias, E., Gampa, K., Golji, J., Kao, I., Li, L., Megel, P., Perkins, T. A., Ramadan, N., Ruddy, D. A., Silver, S. J., Sovath, S., Stump, M., Weber, O., Widmer, R., Yu, J., Yu, K., Yue, Y., Abramowski, D., Ackley, E., Barrett, R., Berger, J., Bernard, J. L., Billig, R., Brachmann, S. M., Buxton, F., Caothien, R., Caushi, J. X., Chung, F. S., Cortes-Cros, M., deBeaumont, R. S., Delaunay, C., Desplat, A., Duong, W., Dwoske, D. A., Eldridge, R. S., Farsidjani, A., Feng, F., Feng, J., Flemming, D., Forrester, W., Galli, G. G., Gao, Z., Gauter, F., Gibaja, V., Haas, K., Hattenberger, M., Hood, T., Hurov, K. E., Jagani, Z., Jenal, M., Johnson, J. A., Jones, M. D., Kapoor, A., Korn, J., Liu, J., Liu, Q., Liu, S., Liu, Y., Loo, A. T., Macchi, K. J., Martin, T., McAllister, G., Meyer, A., Molle, S., Pagliarini, R. A., Phadke, T., Repko, B., Schouwey, T., Shanahan, F., Shen, Q., Stamm, C., Stephan, C., Stucke, V. M., Tiedt, R., Varadarajan, M., Venkatesan, K., Vitari, A. C., Wallroth, M., Weiler, J., Zhang, J., Mickanin, C., Myer, V. E., Porter, J. A., Lai, A., Bitter, H., Lees, E., Keen, N., Kauffmann, A., Stegmeier, F., Hofmann, F., Schmelzle, T., and Sellers, W. R. (2017) Project DRIVE: A Compendium of Cancer Dependencies and Synthetic Lethal Relationships Uncovered by Large-Scale, Deep RNAi Screening. Cell 170, 577–592 e510

56. Ianevski, A., Giri, A. K., and Aittokallio, T. (2020) SynergyFinder 2.0: visual analytics of multi-drug combination synergies. Nucleic Acids Res 48, W488–W493

57. Bliss, C. I. (1939) The toxicity of poisons applied jointly. Annals of Applied Biology

58. Potdar, S., Ianevski, A., Mpindi, J. P., Bychkov, D., Fiere, C., Ianevski, P., Yadav, B., Wennerberg, K., Aittokallio, T., Kallioniemi, O., Saarela, J., and Ostling, P. (2020) Breeze: an integrated quality control and data analysis application for high-throughput drug screening. Bioinformatics 36, 3602–3604

59. Sinijarv, H., Wu, S., Ivan, T., Laasfeld, T., Viht, K., and Uri, A. (2017) Binding assay for characterization of protein kinase inhibitors possessing sub-picomolar to sub-millimolar affinity. Anal Biochem 531, 67–77

60. Meissner, P., Pick, H., Kulangara, A., Chatellard, P., Friedrich, K., and Wurm, F. M. (2001) Transient gene expression: recombinant protein production with suspension-adapted HEK293-EBNA cells. Biotechnol Bioeng 75, 197–203

